# Processing and analyzing high-throughput microfluidic enzymology data: a practical guide to rate fitting and quality control

**DOI:** 10.64898/2026.09.21.753063

**Authors:** Nicholas J. Freitas, Jonathan S. Zhang, Duncan F. Muir, Hayden S. Saunders, Dylan Aidlen, Margaux M. Pinney

## Abstract

High-throughput enzymology enables quantitative characterization of enzyme function across hundreds to thousands of sequence variants and experimental conditions. Nevertheless, the scale and complexity of these datasets create substantial challenges for analysis and quality control. High-Throughput Microfluidic Enzyme Kinetics (HT-MEK), for example, generates large microscopy datasets that must pass through multiple analytical stages, including image processing, initial-rate fitting, and kinetic modeling. Choices or errors made at any stage can propagate into the final kinetic parameters without being evident from fit statistics alone. Here, we provide a broadly applicable guide for analyzing high-throughput enzymology data from the HT-MEK platform, using Michaelis-Menten kinetics as a representative example. We first outline the conceptual workflow from raw fluorescence measurements to estimates of *k*_cat_, *K*_M_, and *k*_cat_/*K*_M_. We then discuss common experimental and analytical failure modes and present a framework for deciding when data should be refitted, filtered, qualified, or repeated. Finally, we provide a step-by-step workflow using *Mercury*, an open-source Python framework that integrates scalable HT-MEK data processing with traceable quality control and diagnostic visualization. This workflow preserves the connection between reported parameters and their underlying measurements and can be adapted to other kinetic models and biochemical assays.

## 1. Introduction

High-throughput enzymology, exemplified by the <u>H</u>igh-<u>T</u>hroughput <u>M</u>icrofluidic <u>E</u>nzyme <u>K</u>inetics (HT-MEK) platform (Markin et al., 2021, 2023; Muir et al., 2025), enables the parallel quantitative measurement of enzyme kinetic parameters across hundreds to thousands of sequence variants and conditions, with replicates. These experiments have revealed the functional architecture of alkaline phosphatase(Markin et al., 2021, 2023), mapped kinetic diversity across evolution and disrupted a widely held model about activity-temperature relationship in adenylate kinase (Muir et al., 2025), and revealed catalytic, specificity, and inhibitory strategies of the SARS-CoV-2 main protease (Aidlen et al., 2026).

HT-MEK belongs to a broader family of massively parallel methods for mapping protein sequence to molecular phenotype. These include pooled deep-mutational-scanning assays typically based on growth selection or reporter-guided enrichment and sorting (Fowler et al., 2010; Fowler & Fields, 2014; Rollins et al., 2019), display-based measurements of binding affinity (Adams et al., 2016; Aditham et al., 2021), multiplexed measurements of protein abundance and folding stability (Amorosi et al., 2021; Matreyek et al., 2018; Tsuboyama et al., 2023), and arrayed or flow-cell platforms for direct biochemical measurements of binding and catalysis (Layton et al., 2019). Many of these approaches achieve scale by reducing each variant to an enrichment score, an endpoint signal, or a small set of summary phenotypes. HT-MEK occupies a complementary niche: by independently estimating enzyme concentration and monitoring reaction progress across substrate concentrations and replicates, it enables estimation of catalytic parameters for each enzyme variant. These measurements more closely resemble classical biochemical characterization and provide more direct access to mechanistic variation across sequence space.

Despite their mechanistic power, these experiments pose a substantial but information-rich analytical challenge. Raw data can comprise up to hundreds of gigabytes of microscopy images, typically encoding 10^5^–10^7^ experimental datapoints that are then fit to obtain kinetic and thermodynamic constants. This expansive raw dataset can be examined at multiple levels, from individual images and reaction-progress curves to spatial patterns, signal-to-concentration relationships, replicate consistency, and kinetic-model fits. Realizing this potential requires a multistep analysis in which image processing, background subtraction, fitting-window selection, rate estimation, replicate handling, and quality-control decisions remain explicit and inspectable. Because manual inspection and spreadsheet-based analysis are impractical at this scale, an effective workflow must combine automation with diagnostic visualization, using the complete scope of the data to reveal experimental and analytical problems rather than allowing them to remain hidden.

This guide will serve as both a primer for scientists analyzing HT-MEK datasets for the first time and a reference for regular HT-MEK users. We focus on the analysis that begins once the assay, imaging scheme, and kinetic model have been selected: image processing, calibration, rate fitting, quality control, parameter estimation, and reporting. We do not cover system selection, assay design, imaging setup, or formulation of an appropriate kinetic model in detail. Instead, this guide begins once these decisions have been made and an experimental dataset is in hand.

The overall analysis framework builds directly on the pipeline developed in the Fordyce laboratory and described in Markin et al. (2021). We retained much of the underlying analytical logic and sequence, with some laboratory-specific differences, while substantially redeveloping and extending the implementation as *Mercury*, an open-source Python framework. *Mercury* streamlines the workflow, provides diagnostic visualization throughout data processing and analysis, facilitates the detection and correction of analytical errors, and improves usability, adaptability, and maintainability. Other laboratories may reasonably adopt different choices at individual steps depending on their experimental systems and analytical priorities; our goal is to make our own choices and their rationale explicit and provide a flexible framework for implementing changes.

For the purpose of this guide, we will walk through the analysis of Michaelis-Menten kinetics to fit the following constants: catalytic turnover (*k*_cat_), Michaelis Constant (*K*_M_), and catalytic efficiency (*k*_cat_/*K*_M_). While the methodology described herein is easily extended to other kinetic models, fitting Michaelis-Menten parameters provides a straightforward example that illustrates the major steps in data processing, rate fitting, and quality control.

Section 2 develops the measurement and modeling logic that connects raw fluorescence to kinetic parameters. Section 3 describes common ways that real high-throughput datasets deviate from this ideal and provides a decision framework for their diagnosis and resolution. Finally, Section 4 provides a walk-through of the analysis workflow implemented in *Mercury*, a software package for scalable analysis of high-throughput enzyme kinetics, including data organization, calibration, initial-rate fitting, quality-control filtering, Michaelis-Menten fitting, and export of interpretable results (**Fig. 1**).

**Figure 1.**
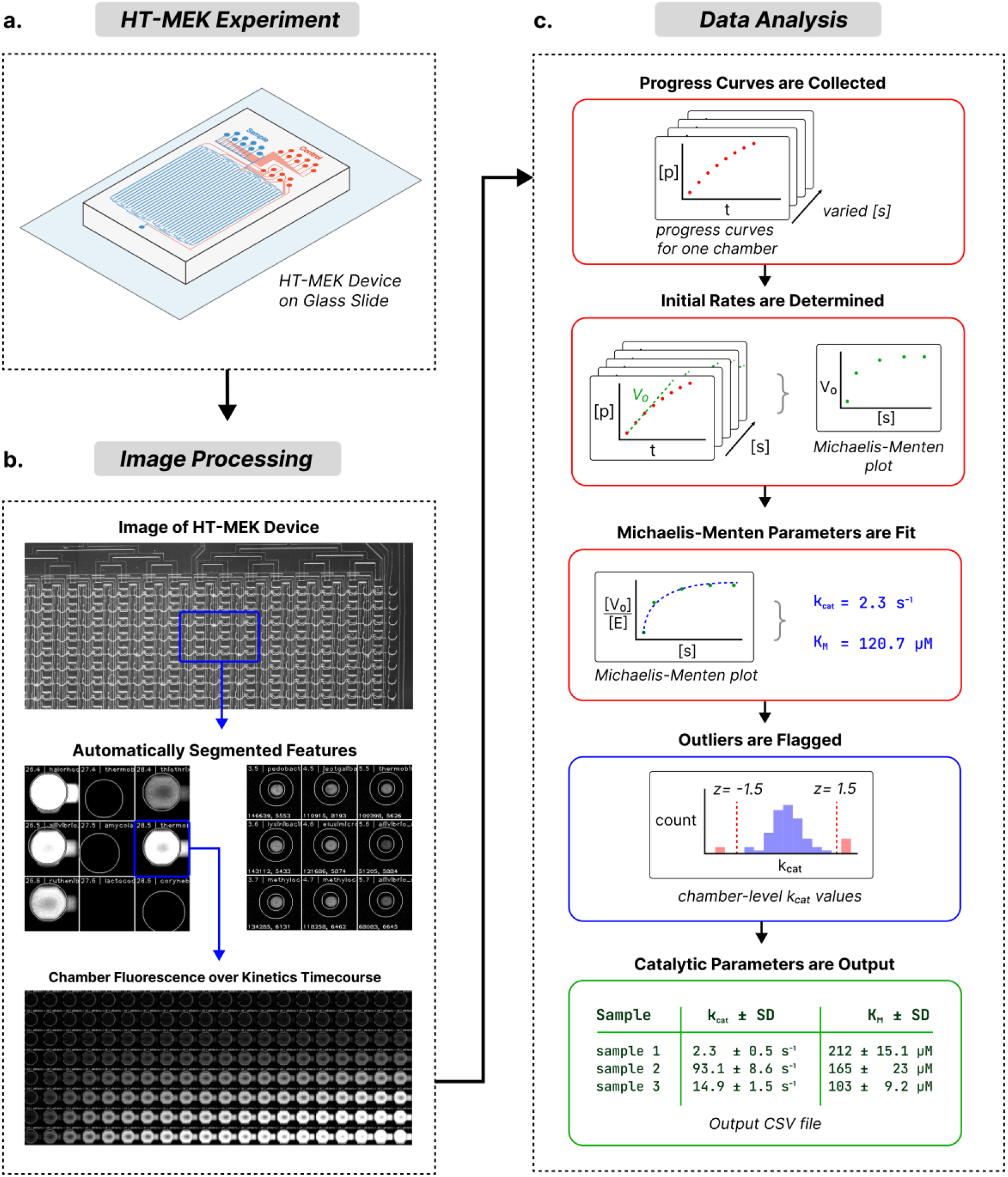
End-to-end analysis of HT-MEK data yields Michaelis-Menten parameters. **A.** Product standard curve and enzyme kinetic measurements are taken on an HT-MEK device. **B.** Raw microscope images are merged and segmented to identify fluorescent device features. The aggregate fluorescence for each chamber is used to derive a fluorescence vs. time series at each substrate concentration. **C.** Product standard curves are used to create product vs time series for each chamber, at each substrate concentration. Initial rates are fit to each time-course, and used to construct a Michaelis-Menten curve for each chamber. Kinetic parameters k_cat_ and K_M_ are fit, aggregated among replicates, and outliers are identified for inspection. Final catalytic parameters are output.

## 2. Measurement and modeling framework for HT-MEK

### 2.1. Michaelis-Menten Kinetics

In this guide, we will work through the analysis of Michaelis-Menten kinetics from HT-MEK experimental data. The minimal irreversible mechanism is E + S ⇌ ES → E + P. Under the steady-state approximation and initial-rate conditions, this mechanism yields a hyperbolic dependence of rate on substrate concentration. More complex mechanisms can produce the same empirical form, so the fitted parameters must be interpreted in the context of the mechanism and assay.

We typically monitor product formation, substrate loss, or a stoichiometrically coupled reporter. Restricting the fit to ≤10% of reaction progress keeps substrate concentration close to its initial value and reduces product-dependent effects and reverse reaction.

Under these conditions, the Michaelis-Menten equation is defined as:

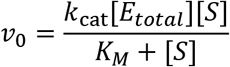

Here, [E]_T_ is total enzyme concentration, *k*_cat_ is the catalytic turnover number, and *K*_M_ is the substrate concentration at which v_0_ (initial velocity) reaches half of *V*_max_. Operationally, *K*_M_ describes the shape of the steady-state rate curve and should not be equated generally with a binding dissociation constant. For the minimal irreversible mechanism above, *K*_M_ = (*k*_cat_ + *k*_off_)/*k*_on_; *K*_M_ approaches the ES dissociation constant only when catalysis is much slower than substrate dissociation. Measuring v_0_ across substrate concentrations allows *V*_max_ and *K*_M_ to be fitted, after which *k*_cat_ = *V*_max_ /[E]_T_.

While we focus on Michaelis-Menten kinetics in this guide, the HT-MEK platform offers flexibility to explore alternate systems, including binding kinetics or inhibition constants (Aidlen et al., 2026; Markin et al., 2021, 2023). In these advanced use cases, the acquisition, merging, and segmentation of microscope images will be nearly identical to the descriptions in this guide. When analyzing data with *Mercury* (**Section 4**), users may specify custom kinetic equations, and the exact workflow will vary based on the desired results. In all cases, carefully evaluate the mathematical assumptions of your chosen kinetics model, and use that information to guide experimental data acquisition, model fitting, calibration, and quality control.

### 2.2. How HT-MEK instantiates this experiment

HT-MEK implements this basic enzyme kinetics experiment at high throughput. In a typical device, many unique enzyme variants and their replicates are expressed, captured, and purified in 1,792 individual reaction chambers (Aidlen et al., 2026; Markin et al., 2021, 2023; Muir et al., 2025) (**Fig. 3a-c**). Pneumatically actuated valves direct fluid flow and isolate each chamber, while a circular “button” valve controls fluid exposure to a defined surface region of each chamber typically in the center of each reaction chamber during the experimental workflow (**Fig. 3b).** Enzyme concentration is estimated from a fluorescent tag, most often, eGFP, using a separately established calibration curve, and reaction progress is monitored through changes in concentration over time of a fluorescent product, substrate, or coupled fluorescent readout.

To acquire an image of the full device, a grid of sub-images, often 5×5 under our typical imaging conditions (**Section 4**), is taken, which are then merged (or “stitched”) together into a contiguous image. Physical features in the microfluidics device, such as the reaction chambers, are identified via image segmentation. The summed product fluorescence of these segmented features is monitored over time.

During analysis, fluorescence measurements must be converted to product concentration using standard curves; for coupled or substrate-loss assays, the calibrated reporter-equivalent concentration must be defined, along with its sign and stoichiometric relationship to reaction turnover. Next, the product concentration traces must be fit over an appropriate time window to estimate initial linear rates (*v_0_*). Finally, those rates must be combined across substrate concentrations and replicates to estimate kinetic parameters. This analysis must be performed in parallel for all 1792 reaction chambers of the device. It must be robust and adaptable to measure a large dynamic range of initial rates. And because an experiment may have hundreds of samples each with tens of replicates, metadata including chamber identity must be tracked to merge and filter replicates at the end.

### 2.3. From fluorescence signal to product concentration

In a typical HT-MEK experiment, we estimate product formation by measuring the summed pixel intensity of fluorescence within each reaction chamber, reported in relative fluorescence units (RFUs) (**Fig. 1B**). The integrated fluorescence signal, reported here as summed RFU, using our typical microscopy setup (see **Section 4**) depends not only on fluorophore abundance within the segmented region but also on camera exposure time, excitation irradiance, optical-path transmission, detector response, background fluorescence and detector offset, and the area included in the segmentation mask. These confounding factors can vary along two distinct axes: over <u>time</u>, across the frames of a single time course, and over <u>space</u>, across the field of view of the device.

Temporal variation can arise when imaging conditions change over the course of an experiment, such as ambient light changing and being detected by the microscope. These changes can most easily be controlled with careful precautions at acquisition time.

Spatial variation, however, arises because optical and detection properties differ across the device, and this can be corrected during the data analysis stage. Illumination and background fluorescence vary from chamber to chamber, properties which are minimized by taking one or more *background* images without substrate or product in the device and subtracting this image from assay images before analysis. (**Fig. 2B)** This background subtraction greatly reduces additive signals arising from stray light, device autofluorescence, and stationary debris. A separate confounding effect is spatial nonuniformity across the microscope field of view. Nonuniform excitation irradiance and vignetting in the detection path can cause the same fluorophore concentration to produce a stronger measured signal near the center of the field than at its edges. We account for this position-dependent response by fitting a separate product standard curve for each chamber and using that curve to convert the measured fluorescence signal into product concentration for that chamber (Fig. 2D).

**Figure 2.**
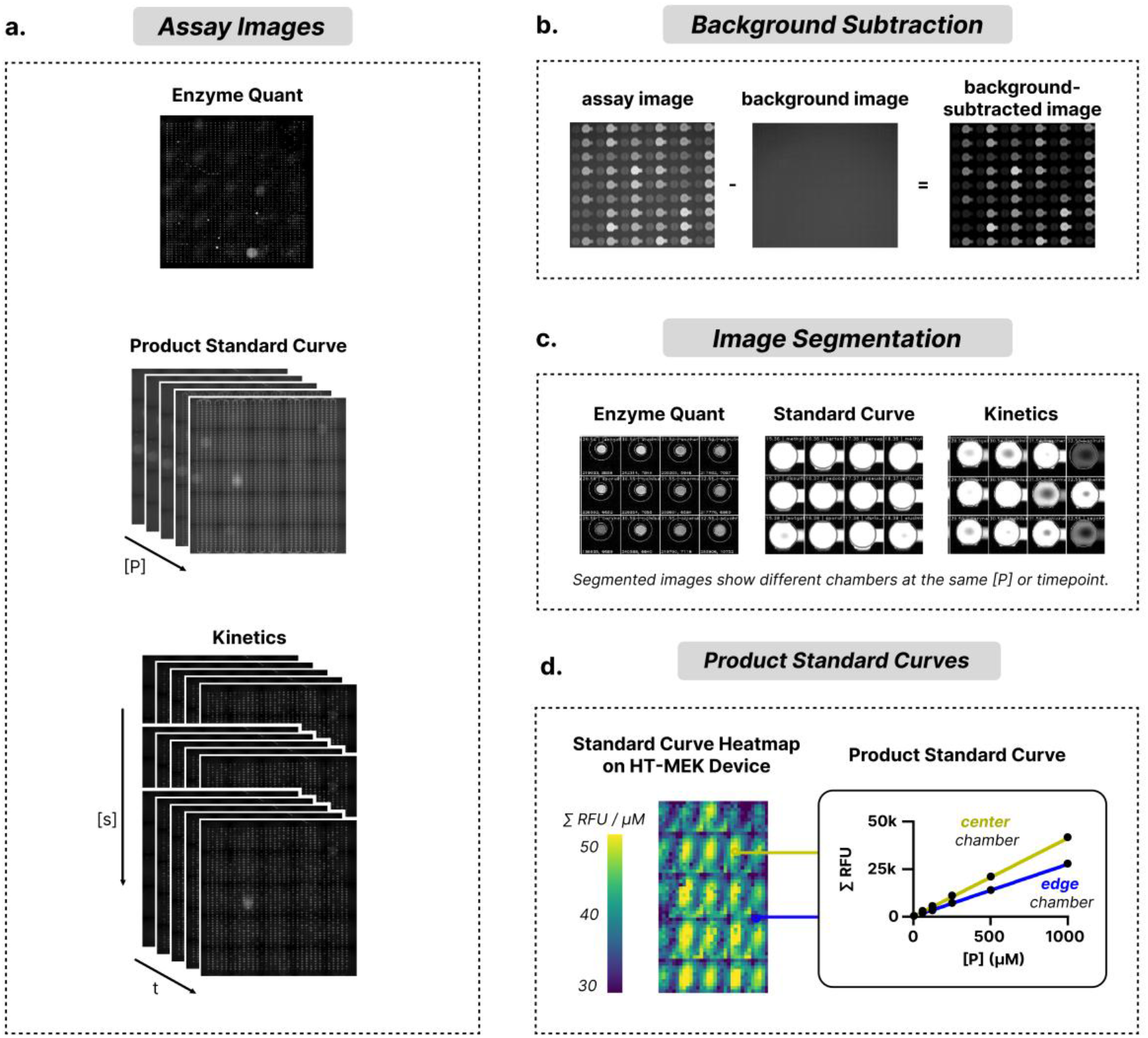
Raw images are processed to yield initial rate fits. **A.** Experimental data comprises product standard images, enzyme kinetics time-courses, and a single image to quantify eGFP-tagged enzyme concentration. Additionally, *background images* are taken (not shown). **B.** The fluorescence values from background images are subtracted from corresponding kinetics or standard curve images. This reduces erroneous fluorescence from sources including ambient light or debris. **C.** Images are automatically segmented to determine the location of relevant device features, so aggregate fluorescence can be calculated. **D.** The optics within the microscope camera and light-path cause lower sensitivity to fluorescence at the edge of images. This effect is counteracted by measuring a unique product standard curve for each well. Different standard curve slopes capture the effect of optical vignetting.

As is typical of product standard curves, a series of known product concentrations is introduced into the device and imaged under the same conditions as the kinetic assay. For each chamber, the relationship between product concentration and summed chamber RFUs can then be fit to generate a standard curve. If the signal is linear over the relevant range, the slope of the standard curve gives the ratio of chamber fluorescence to product concentration for that chamber. During kinetic analysis (**Section 2.4**), the fluorescence trace of each chamber is converted to product concentration using only that chamber’s own standard curve. In this way, the per-chamber product standard curves counteract the effects of spatial variance in camera sensor sensitivity, local illumination differences, or background fluorescence.

### 2.4. From a change in [Product] to initial rates

Once product concentration has been inferred for each chamber and time point, the next goal is to determine the initial reaction rate, *v_0_*. In practice, *v_0_* is estimated by fitting a line to an early portion of the product-versus-time trace. Under ideal initial-rate conditions, this early region is approximately linear. The slope of this linear fit (with dimensionality of product concentration / time) is then used as the initial rate for that chamber and substrate concentration (**Fig. 1E**).

The choice of fitting window is therefore a central analysis decision. If the window is too long, it will include curvature caused by substrate depletion, resulting in a poor linear fit. If the window is too short, the rate estimate may be dominated by measurement noise. Choosing a fit window that comprises ≤ 10% of the reaction is a sensible start (Srinivasan, 2022), but one must examine and modify their fit window according to the needs of each dataset. In **Section 3.5-3.6**, we discuss common sources of error, worthwhile considerations, and diagnostics during initial-rate fitting.

In some experimental systems, the non-enzymatic rate of the reaction will be non-negligible, due to high intrinsic reaction rate in solution. If uncorrected, a background rate will systematically bias downstream *k*_cat_ and *K*_M_ fits. Additionally, a background rate may obscure the activity of weakly active enzyme variants and set a lower limit on the catalytic rates that can be measured. For this reason, it is preferable to calculate and subtract the background rate from the measured sample initial rates. This can be accomplished in practice by including several *no-enzyme* (or buffer-control) chambers in an HT-MEK experiment. To conservatively estimate the background rate, we use the median of the initial rates of the *no-enzyme* chambers for each substrate concentration. This rate is then subtracted from the initial rates of all other samples at the corresponding substrate concentration.

### 2.5. From initial rates to Michaelis-Menten parameters

For each HT-MEK device chamber, the parameters *k_cat_* and *K_M_* are calculated by fitting initial rates across substrate concentrations to the enzyme-normalized Michaelis-Menten equation:

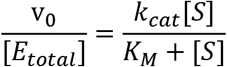

To estimate *k*_cat_ and *K*_M_ robustly, we must sample sufficient points along this curve. Data collected at high substrate concentrations, where the rate approaches a plateau, constrain *k*_cat_, whereas data collected in the low-substrate, approximately linear regime, constrain *k*_cat_*/K*_M_. Generally, we aim to assay substrate concentrations that are one order of magnitude above and below *K*_M_ (0.1×-10× *K*_M_). This range may be inaccessible due to limitations of substrate solubility, or when *K*_M_ varies greatly for different enzyme variants measured in the same HT-MEK experiment; in the former case, we may instead estimate the specificity constant *k*_cat_*/K*_M_ from the linear region of the *v*_0_ vs [S] curve. In the latter case, the range of [S] measured in an experiment may need to be expanded beyond our ± 10x “rule-of-thumb”

After kinetic parameters have been estimated, replicate measurements are evaluated for technical failures and consistency before being combined. For each variant, retained replicates are combined to obtain summary parameter estimates and associated uncertainty. **Section 3.7** describes the diagnostics used to assess these parameter estimates and fit quality, identify technical failures, and determine whether measurements should be retained, qualified, refit, or repeated.

## 3. Identifying and addressing common challenges

The workflow described above depends on a chain of testable assumptions: chambers are correctly identified, fluorescence values are reliable, enzyme and product concentrations are detectable, standard curves are linear, initial rates are derived from the initial linear regime, and the measurements span a sufficient range of substrate concentrations to constrain the

Michaelis-Menten parameters. In high-throughput experiments, some fraction of measurements will inevitably violate these assumptions locally, spatially, or globally.

An analysis pipeline must therefore do more than return best-fit parameters. It should connect each experiment to the underlying measurements, reveal departures from the starting experimental or analytical assumptions, distinguish experimental failures and technical artifacts, and decide whether a dataset should be retained, re-fit, filtered, or repeated. The goal of quality control is to identify which assumption in the analysis chain has failed, to choose a response appropriate to the scale and cause of that failure. When exclusion is warranted, the applicable criterion and rationale should be documented and reported.

In this section, we discuss common ways that HT-MEK datasets deviate from the ideal workflow. Rather than presenting these issues as an exhaustive list of possible artifacts, we organize them by the stage of analysis at which they become visible. For each class of problem, the reader should ask three questions:

1) What diagnostic plot, image, or metric reveals the problem?
2) Is the likely cause experimental, analytical, or model based?
3) Can the issue be corrected by changing analysis parameters, handled by filtering, reported with qualification, or does it require repeating the experiment?

### 3.1. Diagnosing Experimental and Analytical Failure Patterns

When an unexpected pattern appears in an HT-MEK dataset, it is important to determine both what is affected and how broadly the issue extends Within a given chip, the spatial distribution of measurements can help distinguish isolated chamber-level abnormalities, regional patterns, and device-wide problems. We therefore recommend mapping key measurements and fit diagnostics onto chamber locations and examining these heatmaps alongside raw or summary images, controls, and replicate plots (see Sections 4.3–4.6).

A <u>local failure</u> affects only a small number of chambers. Examples include dust, aggregates in a chamber, poorly segmented chambers, unusually low enzyme expression, or isolated chambers with poor standard curve fits. (**Fig. 3A**) Local failures often appear as isolated outliers in spatial plots of the microfluidics device, summary images, or replicate plots. These are the best-case failures; if the rest of the dataset is well-behaved, these measurements can often be filtered without compromising the entire experiment.

**Figure 3.**
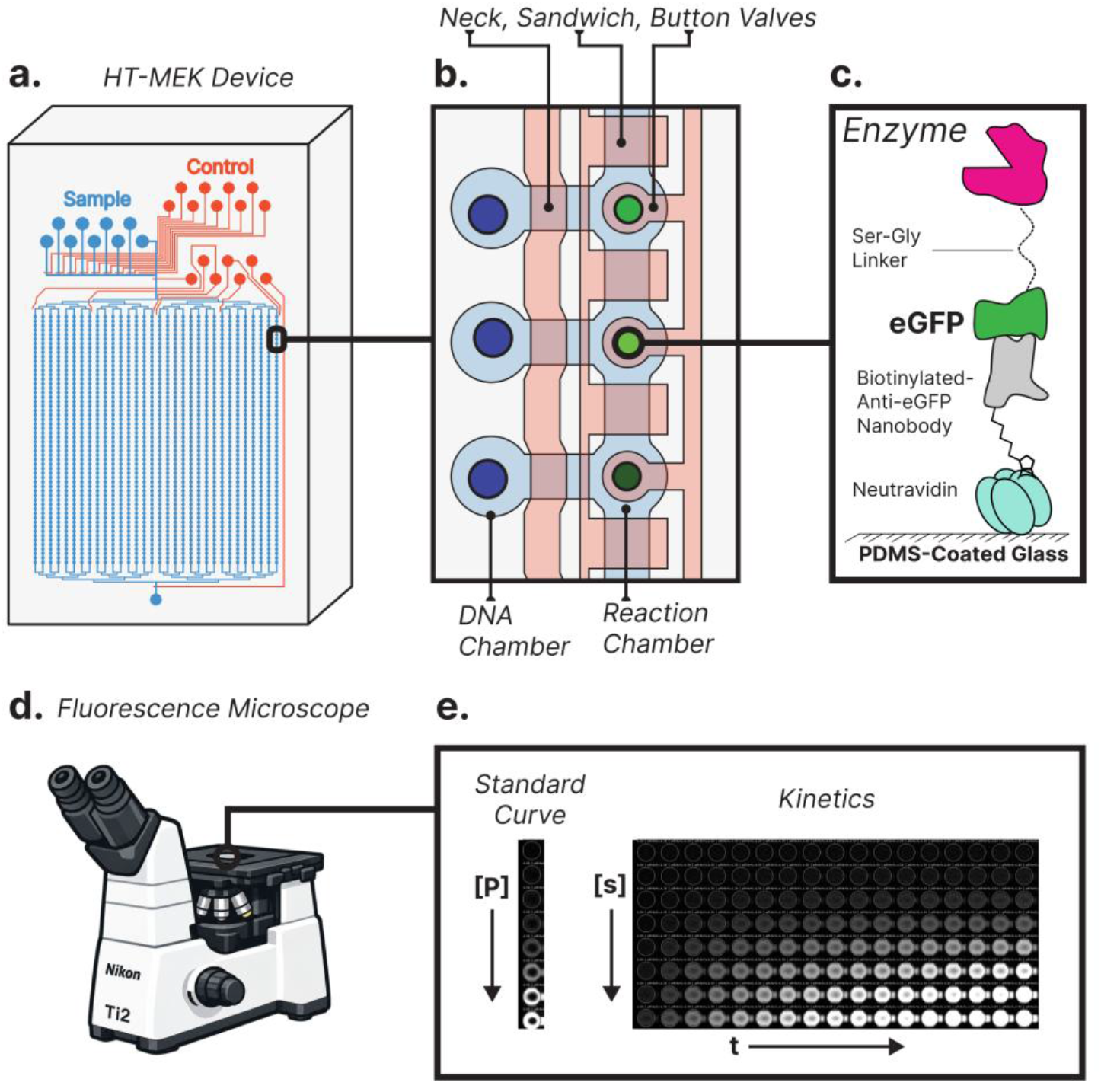
Schematic of the HT-MEK platform. **A.** The HT-MEK microfluidics device contains 1,792 individual reaction chambers. **B.** During an HT-MEK assay, chambers are isolated from each other and enzymes are expressed. **C.** eGFP-tagged enzymes are localized within the reaction chamber using an anti-eGFP nanobody. **D.** Fluorescence in each chamber is tracked using a Nikon Ti2 Fluorescence Microscope. **E.** Product standard curve images are captured for each chamber. Then, kinetic activity is tracked via product fluorescence.

**Figure 4.**
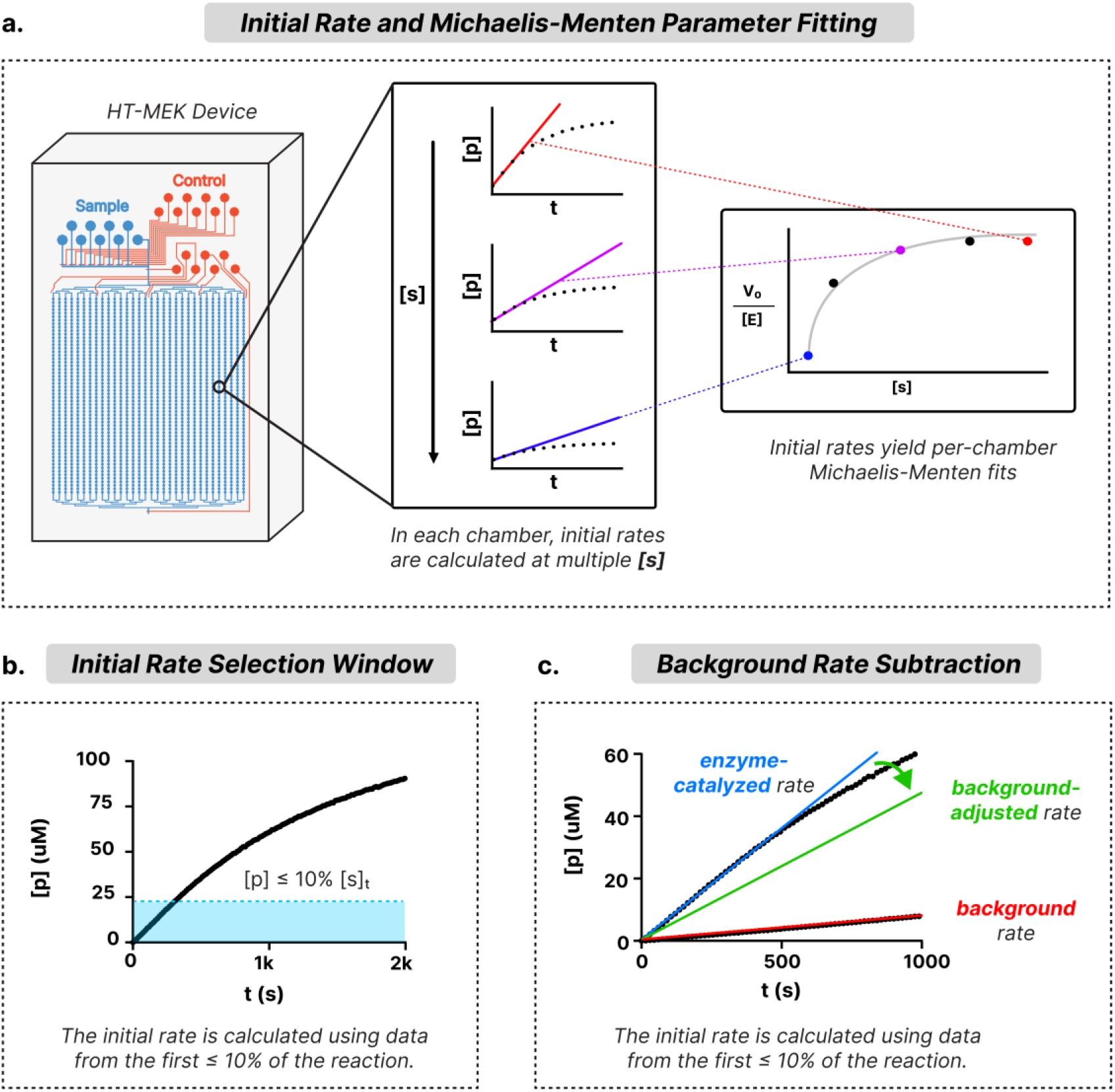
Workflow for initial-rate determination and Michaelis–Menten fitting in HT-MEK. **A**. For each chamber, initial rates are calculated at varying substrate concentrations. These are used to plot per-chamber Michaelis-Menten curves for subsequent fitting of *k*_cat_ and *K*_M_. **B.** Initial rate must be fit to the early, linear portion of the reaction. For this reason, data from no later than the first 10% of the reaction should be used. **C.** Some reactions have considerable non-enzymatic background rates. To account for this, the initial rate of the reaction is calculated at each substrate concentration using *no-enzyme* chambers. This background rate is subtracted from the calculated initial rates before further analysis.

A <u>spatially patterned failure</u> affects a region of the device, such as a row, column, edge, tile, or gradient across the chip. These patterns often indicate device-level or imaging-level artifacts: poor flow through a set of channels, valve actuation errors, uneven illumination, or image stitching artifacts. To avoid errors in which spatially patterned failures affect many replicates of the same sample, skewing results, HT-MEK experiments randomize the location of replicates across the device. In cases of spatially patterned failures, the appropriate response is often filtering out affected regions - for example, an entire column that experienced low substrate concentration due to poor reagent flow.

A <u>global failure</u> affects most or all the dataset. Examples include poor enzyme expression across the device, product fluorescence that saturates the camera detection limit, product fluorescence outside the calibration range, failed substrate loading, air ingress into the device, or uniformly weak product signal. Global failures are less likely to be rescued by filtering, which does not fix the underlying measurement issue. These are worst-case results, as they often require repeating the experiment or changing the experimental design.

Finally, a <u>model-level failure</u> occurs when the measurements themselves are valid, but the model or fitting strategy does not support the expected interpretation. Examples include product accumulation traces that are real but not linear over the selected fitting window, substrate titrations that do not constrain both *K*_M_ and *k*_cat_, or rate data that describe a background process rather than the enzyme-catalyzed reaction. In these cases, the appropriate response is not always to repeat the experiment. Instead, one may need to change the rate-fitting window, report only a subset of parameters, or qualify the interpretation of the fitted values. In the worst case, the experiment may need to be repeated with different substrate, expression, or imaging conditions to allow for robust characterization of enzyme kinetics.

### 3.2. Choosing the response: refit, filter, qualify, or repeat

This distinction between local, spatial, global, and model-level failures provides a triage framework. Depending on the breadth and intensity of the issue, the researcher should select one of four potential interventions: refit, filter, qualify, or repeat (**Table 1**).

**Table 1:**
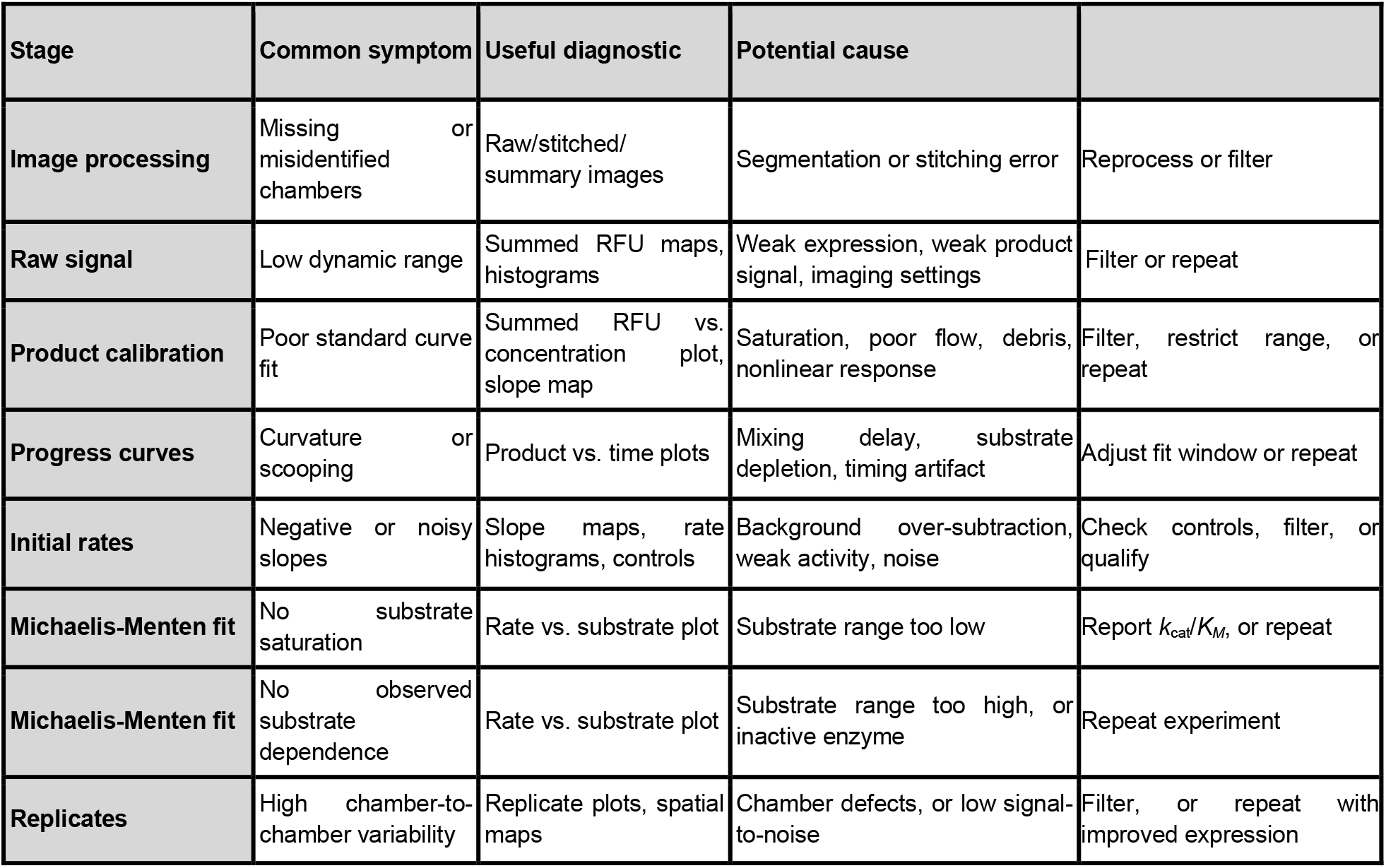
Common symptoms, diagnostics, and responses.

| Stage | Common symptom | Useful diagnostic | Potential cause |  |
| --- | --- | --- | --- | --- |
| Image processing | Missing or misidentified chambers | Raw/stitched/summary images | Segmentation or stitching error | Reprocess or filter |
| Raw signal | Low dynamic range | Summed RFU maps, histograms | Weak expression, weak product signal, imaging settings | Filter or repeat |
| Product calibration | Poor standard curve fit | Summed RFU vs. concentration plot, slope map | Saturation, poor flow, debris, nonlinear response | Filter, restrict range, or repeat |
| Progress curves | Curvature or scooping | Product vs. time plots | Mixing delay, substrate depletion, timing artifact | Adjust fit window or repeat |
| Initial rates | Negative or noisy slopes | Slope maps, rate histograms, controls | Background over-subtraction, weak activity, noise | Check controls, filter, or qualify |
| Michaelis-Menten fit | No substrate saturation | Rate vs. substrate plot | Substrate range too low | Report $k_{cat}/K_M$, or repeat |
| Michaelis-Menten fit | No observed substrate dependence | Rate vs. substrate plot | Substrate range too high, or inactive enzyme | Repeat experiment |
| Replicates | High chamber-to-chamber variability | Replicate plots, spatial maps | Chamber defects, or low signal-to-noise | Filter, or repeat with improved expression |

<u>Refit</u> initial rates or Michaelis-Menten parameters when the underlying data are valid, but the analysis choice was inappropriate. Examples include an initial-rate window that includes late-time curvature, an early time point affected by mixing, poor image segmentation, or a background rate estimate that was applied incorrectly. Many analytical failures can be resolved through altering processing parameters. Any changes should be scientifically justified, applied consistently to comparable measurements, and documented.

<u>Filter</u> your data when the problem is localized, and the remaining data still supports the downstream analysis. Examples include isolated chambers with poor segmentation, debris, or low enzyme expression. Filtering should remove measurements that violate analysis assumptions, but it should not be used to force a desired result. To this end, filtering should be based on interpretable criteria, applied consistently, and recorded for reproducibility.

<u>Qualify</u> your results when the data quality supports some conclusions but not others. For example, a substrate titration that remains in the low-substrate regime may support *k*_cat_*/K*_M_ estimation but not independent estimates of *K*_M_ and *k*_cat_. Or, certain enzyme variants may have catalytic parameters that are unconstrained by the data. In these cases, the result should be reported with appropriate caveats rather than treated as either fully valid or fully failed.

<u>Repeat</u> an experiment when the failure is global, upstream, or analytically uncorrectable. Examples include failed substrate loading, widespread lack of enzyme expression, or time courses that do not capture an initial-rate regime. Initial, small-scale experiments to validate experimental conditions can reduce the liklihood of this outcome.

### 3.3. Failures in Image Processing: Acquisition, Stitching, and Conversion

Before interpreting rate fits or kinetic parameters, one must establish that the raw measurements are physically meaningful. HT-MEK images must be stitched, background-subtracted, segmented into chambers, and converted into fluorescence values (**Fig. 1**); failures at any of these stages propagate through the remainder of the analysis. Dust, debris, scratches, or optical defects may obscure chambers or create spurious signal (**Fig. 5A,B**), while poor stitching or segmentation may shift, omit, or incorrectly identify device features. Low signal, strong background fluorescence, and uneven illumination can further limit the dynamic range available for detecting enzyme or product fluorescence.

**Figure 5:**
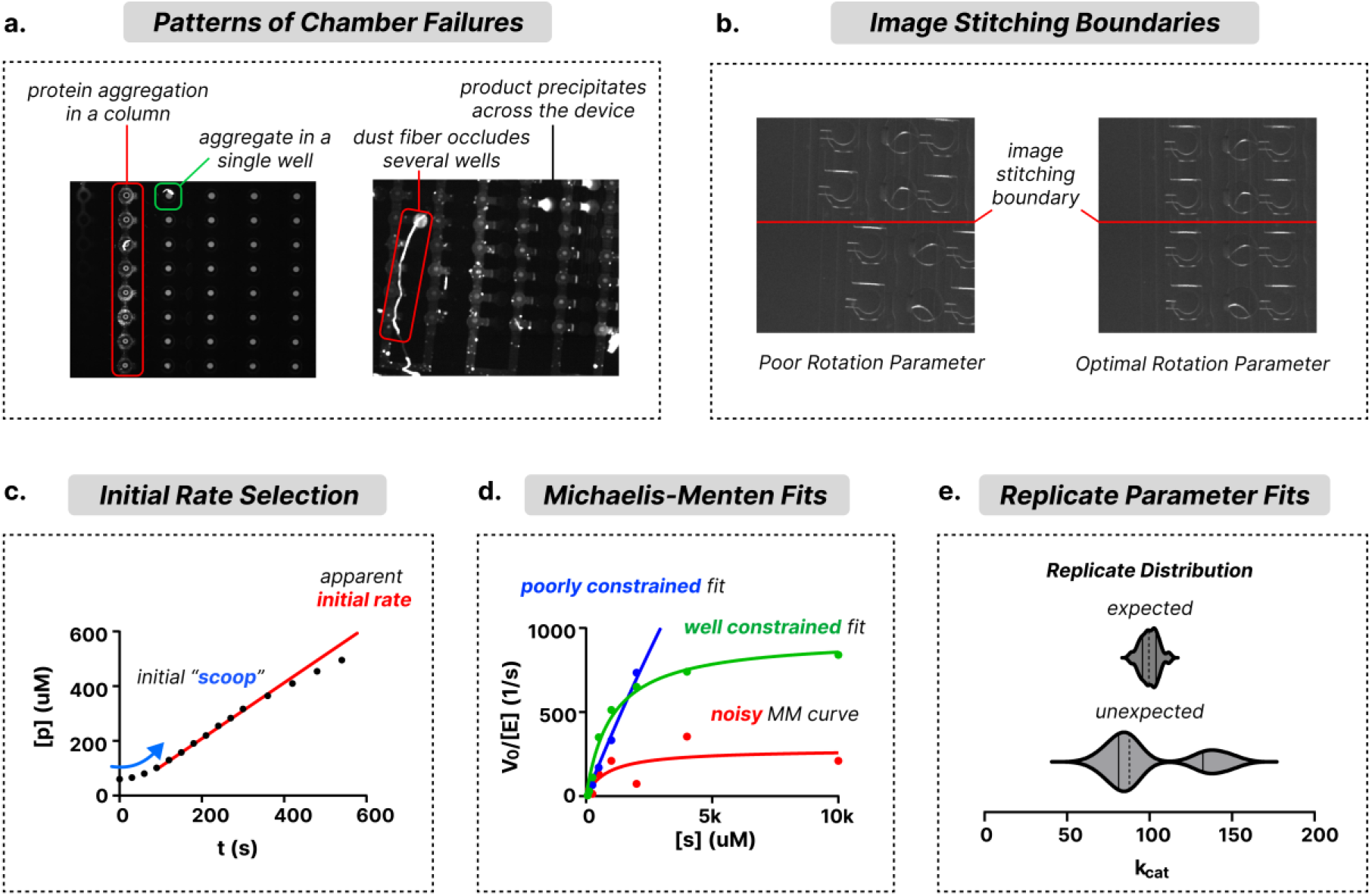
Identifying experimental and analytical failure modes in HT-MEK datasets. **A.** Experimental failures may occur locally, such as aggregates in a single well. They may be spatially patterned, due to protein aggregation or dust affecting groups of wells. Or, they may be global failures, due to experimental conditions that affect the entire device. **B.** Errors during sub-image stitching will affect downstream analysis. Using the optimal image rotation parameter ensures there are no discontinuities across image boundaries. **C.** The initial rate fitting window must be chosen to capture the early, linear portion of the reaction, while avoiding artifacts like “scooping”, where the reaction begins slowly due to incomplete mixing of reactants. **D.** To determine Michaelis-Menten parameters properly, initial-rate measurements must be taken across a sufficiently wide range above and below *K*_M_; otherwise, fits will be poorly constrained. Noisy data will confound Michaelis-Menten curve fitting. **E.** Replicate samples across multiple chambers are compared to assess the quality of fit parameters. Unexpected parameter distributions across replicates (for example, bimodal distributions) may indicate experimental or analysis issues, such as mislabeled chambers, and should prompt manual review.

**Figure 6:**
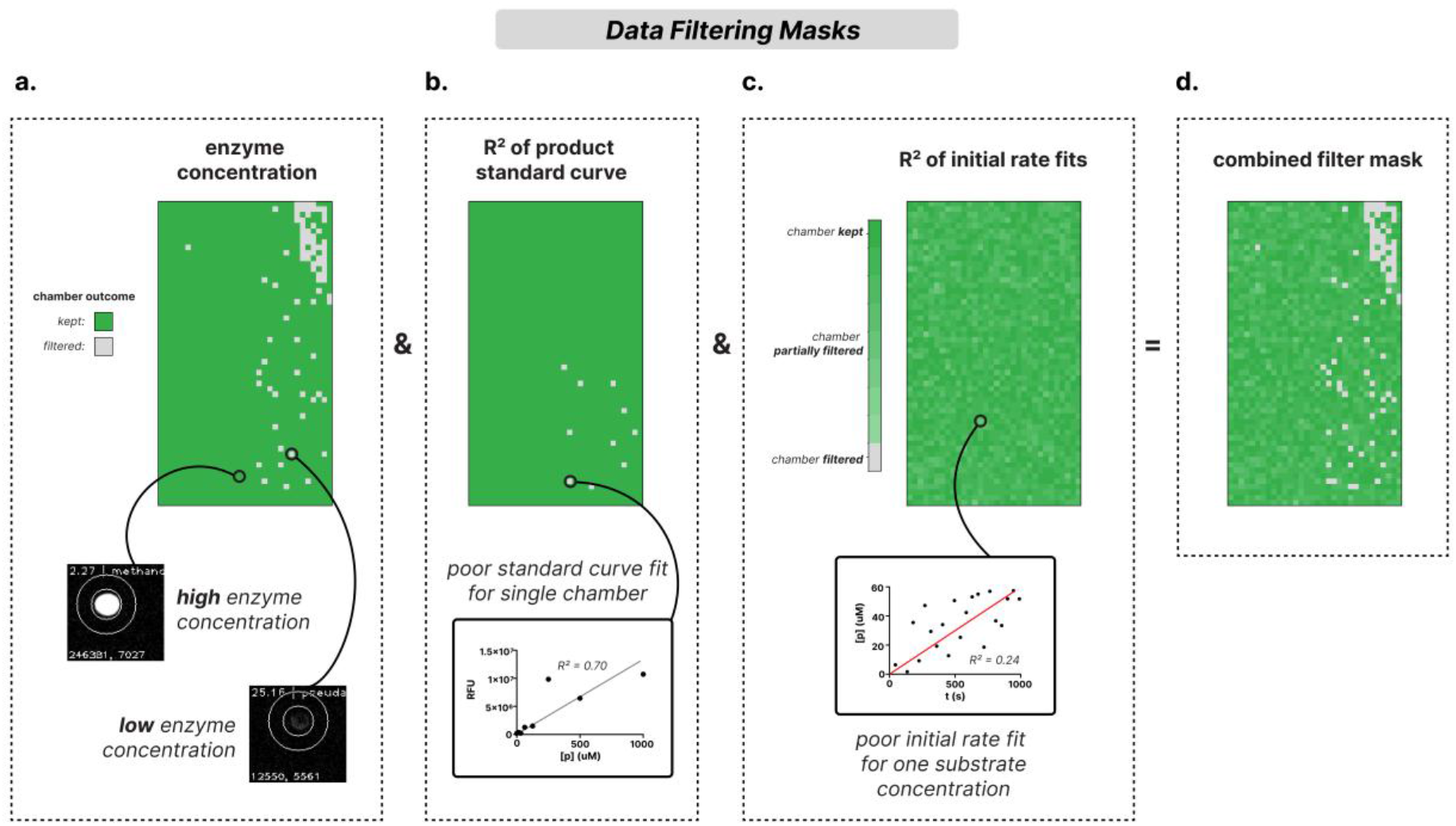
Filtering masks are generated for all chambers and initial rate fits. The reliability of fit parameters is determined for each chamber and substrate concentration through a series of quality filters. Boolean masks are constructed to filter out poor-quality data before downstream analysis. The filtering criteria include minimum enzyme concentration (**A**), minimum R^2^ for per-chamber product standard curves (**B**), and minimum R^2^ for fit initial rates per substrate concentration (**C**). Depending on the criterion, filters may flag either an entire chamber (e.g., insufficient enzyme concentration) or individual initial-rate fits within that chamber (e.g., poor R^2^).Additional filtering criteria (not pictured) include requiring initial rates to exceed the measured reaction background and excluding chambers with insufficient unfiltered substrate concentrations for Michaelis-Menten fitting. These masks are combined via the Boolean *AND* operation (&) to yield a final mask which is applied to the data (**D**).

These problems are most readily diagnosed by inspecting raw and processed images, segmentation summaries, and spatial maps of chamber-level fluorescence. **(Fig 2A,C,D)** Isolated outliers generally indicate local defects, whereas rows, columns, tile-shaped discontinuities, gradients, or edge effects suggest flow, valve, stitching, illumination, or focus problems. Processing errors such as incorrect segmentation or image alignment can often be corrected by reprocessing with different parameters, while a small number of obstructed chambers can be filtered. In contrast, device-wide failures such as absent reagent loading, widespread lack of enzyme signal, or inappropriate imaging settings may leave insufficient information for downstream analysis. Representative diagnostic images are examined in **Section 4.2**.

### 3.4. Standard Curve Failures

Product standard curves convert raw fluorescence into product concentration, so calibration errors introduce systematic bias into all downstream rate estimates. These errors may be difficult to recognize from kinetic fits alone, which can appear well behaved even when the underlying concentration values are incorrect. Common failures include nonlinear or saturated fluorescence responses, calibration signals that are weak relative to background, kinetic measurements that fall outside the calibrated concentration range, and isolated or spatially patterned chambers with poorly determined calibration slopes.

Calibration quality should therefore be assessed before initial-rate fitting using plots of summed RFU versus product concentration and spatial maps of calibration slope and fit quality (**Section 4.3**). Some chamber-to-chamber variation in slope is expected because of differences in excitation intensity and fluorescence-detection efficiency vary across the field of view owing to nonuniform illumination, optical vignetting, and spatial variation in detector response; this variation is one reason that standard curves are fit separately for each chamber. Isolated calibration failures can generally be filtered out, whereas systematic nonlinearity may require restricting the usable concentration range, adjusting the imaging exposure, or repeating the calibration. Kinetic measurements outside the validated calibration range should not be included in downstream model fitting.

### 3.5. Failures in Fitting Initial Rates

Calculation of an initial reaction rate, *v*_0_, requires a progress curve with an early region that is approximately linear. Curvature, a delayed increase in signal (“scooping”), a substantial signal offset at the first measured time point, or a flat or noisy trace can make the estimated rate sensitive to the selected fitting window. In coupled assays, scooping may reflect the time required for reaction intermediates to accumulate and for the coupling reactions to reach a steady-state reporting rate. Delayed reagent delivery can produce a similar pattern. These features warrant further inspection but do not, by themselves, establish a particular biochemical mechanism or technical failure **(Fig 5C)**

Progress curves should therefore be evaluated using representative product-versus-time plots, spatial maps of fitted rates and fit quality, and comparisons among replicates. Negative or near-zero slopes, poor linear fits, or strong replicate disagreement may indicate an unreliable rate estimate, but these features should be interpreted together with enzyme concentration, calibration quality, background controls, and raw-signal diagnostics. When a replicate contains a clear initial linear regime but poor linear fits, the most common cause is inappropriate time-window selection.

### 3.6. Selecting the Initial-Rate Fitting Window

The choice of fitting window is one of the most consequential decisions in initial-rate analysis, and in a high-throughput experiment, it cannot be made once and applied uniformly. The fitting window must be long enough to average over measurement noise and determine a precise slope, but short enough to remain within the initial linear regime. These requirements compete with one another: a short window may yield a noisy estimate for slow reactions, whereas a long window may include curvature for fast reactions. Because reaction timescales vary with substrate concentration, we assign a separate fitting window to each concentration rather than applying a single global window. For each chamber and condition, the window is additionally truncated when product formation exceeds 10% substrate conversion, a safeguard against substrate depletion.

Window selection should not be based on maximizing the linear-fit R^2^ alone. Short intervals often produce deceptively high R^2^ values despite containing too few points to determine a reliable slope. Automated windows should instead be evaluated using the number and timing of included measurements, visual inspection of progress curves, replicate agreement, and the resulting rate uncertainty. Window adjustment is appropriate only when the trace contains a usable linear regime; it cannot rescue measurements dominated by noise or signal near the detection limit.

Datasets spanning an exceptionally broad range of activities may require additional flexibility, as fast and slow variants can require different windows even at the same substrate concentration. In these cases, variants may be divided into activity subsets, fitted using windows appropriate to their respective timescales, and recombined before Michaelis–Menten fitting. This approach avoids imposing a compromise window that performs poorly for both nearly inactive and highly active variants.

### 3.7. Failures in Fitting Kinetic Parameters

After initial rates have been estimated, the rate-versus-substrate curve is fit to the Michaelis-Menten equation for each chamber. At this stage, each chamber is analyzed independently. Chamber-level fits are evaluated before parameter estimates from replicate chambers containing the same enzyme variant are compared and aggregated. Nevertheless, successful numerical fitting does not ensure that the resulting parameter estimates are well constrained by the data.

Other Michaelis-Menten failures arise from outlier substrate concentrations or inconsistent replicates. A single substrate point may disproportionately determine the fit, especially if few concentrations remain after filtering; this is best viewed in rate-versus-substrate plots. An outlier at one substrate concentration may indicate poor buffer flow, substrate precipitation, imaging artifacts, or a systematic issue with that concentration series.

The final step in analysis is aggregating fit parameters from replicate chambers. Replicate disagreement may reflect chamber-level noise, inaccurate enzyme quantification, sample misassignment, or true biological heterogeneity. To avoid outlier data erroneously biasing aggregated Michaelis-Menten parameters, outlier points can be flagged using a z-score threshold and examined using diagnostic plots and segmented images. The extent of data heterogeneity should be evaluated using histograms of replicate fits, and withheld outliers should be reported. We generally require a minimum of three replicates, each with no fewer than five concentration points, to fit Michaelis-Menten parameters for a sample.

Fit statistics should be interpreted cautiously. R^2^ should not be the sole criterion that a Michaelis-Menten fit is evaluated; a high R^2^ can accompany poorly constrained parameter estimates, whereas a low R^2^ may reflect model mismatch, low signal, or outlier measurements. Parameter uncertainty, replicate-to-replicate variation, and visual agreement between the data and model are all important for interpretation. In **Section 4.7**, we view end-to-end diagnostic plots generated by *Mercury* for this evaluation.

## 4. Michaelis-Menten Analysis with *Mercury*: from raw HT-MEK data to kinetic parameters

Here, we will walk through a representative HT-MEK dataset from raw image-derived measurements through calibration, rate fitting, quality-control filtering, kinetic parameter fitting, aggregation of replicate fits into summary statistics, and export of final parameters and diagnostic plots.

While this tutorial describes Michaelis-Menten analysis, *Mercury* has similar Jupyter notebooks prepared for analyzing binding or inhibition constants. Additionally, the flexible Python API can be used to fit data to user-defined catalytic models. Note that all functions and parameters are described in detail in the provided tutorial notebooks. In this guide, we will reference only the most important function calls.

### 4.1. Protocol Setup

Required Materials:

- Computer with Python (version 3.12 or greater) and Conda installed
- *Mercury,* available at https://github.com/pinneylab/mercury/
- Template Jupyter notebooks and example data: https://github.com/pinneylab/mercury_notebooks/
- (Recommended) ImageJ

Our default image-processing parameters are tuned to the image characteristics produced by the microscopy setup described below. Other imaging configurations may require adjustment of the stitching and image-processing parameters. Please refer to Muir et al. (2025) for a detailed description of the camera and microscope setup used in our experiments.

After an HT-MEK assay, the experimentalist will have a directory containing tens of gigabytes of raw imaging data, along with metadata files that include experimental conditions and microscope/camera capture settings (**Table 2**). The following sections describe the process of converting the raw data into usable images using the Jupyter notebook *Processing.ipynb* (Kluyver et al., 2016) and analyzing these data using *Analysis.ipynb.* These notebooks contain detailed usage instructions; this guide will cover the essential conceptual components needed to get started.

**Table 2:** Required Inputs for analysis in *Mercury*.

| Input | Purpose |
| --- | --- |
| *experiments/* | Top-level directory of experimental data |
| *raw_images/* | Sub-directory of raw images for each experimental condition/timestep |
| *Imaging.csv* | Metadata which stores the *imaging* conditions for each image (e.g. imaging channel, light source, etc.) |
| *Series_index.json* | Metadata which stores the *experimental* conditions for each image series (e.g. time values, substrate concentrations, etc.) |
| *Experiment.log* | Record of valve actuations and image acquisitions during experiment. Used for debugging. |
| *Pinlist.csv* | Table relating (x,y) chamber position to sample identification. |
| *Poslist/d\*.pos* | Directory containing the “.pos” file, which records microscope stage position settings. |

The user should not modify raw experimental data at any point during data processing. <u>Always keep a backup of this data.</u>

### 4.2. Image processing: stitching, background subtraction, and chamber segmentation

The first stage of the *Mercury* workflow converts raw microscopy images into chamber-level fluorescence measurements. In an HT-MEK experiment, the imaging directory contains tiled microscopy images for enzyme quantification, product standard curves, kinetic time courses, and background controls. Each experimental condition or time point is captured as a set of partially overlapping sub-images, which are later stitched into a complete image of the device. The resulting images are then background-subtracted, segmented to identify the individual reaction chambers and “buttons” where the enzyme is pulled down, and fluorescence of the bound eGFP-tag is quantified to produce the RFU tables used in downstream analysis.

Errors introduced during this stage propagate into every subsequent calculation, so care should be taken to ensure proper image processing. For this reason, we will frequently inspect raw and processed images. This must be done with a scientific image viewer, as raw HT-MEK images are stored as 16-bit TIFF files, each image pixel records the measured camera intensity *without display normalization*. As a result, these files may appear nearly black when opened in a standard image viewer, even when a useful fluorescence signal is present. This is expected: the measured signal often occupies only a small fraction of the full 16-bit intensity range. For visual inspection, images should therefore be viewed using a tool with display-range adjustment, such as ImageJ (Schneider et al., 2012). When adjusting or viewing images, <u>do not save them</u> to avoid overwriting experimental data.

#### Step 1: Stitch raw image tiles

Examine a folder under the *raw_images/* directory. Each experimental condition or time point is captured as 25 sub-images. Viewing any individual raw sub-image reveals that it is typically slightly rotated relative to the image axes. This in-plane rotation reflects a small angular offset between the camera sensor andthe microscope-stage coordinate system. *Mercury* therefore estimates the rotation required to align the device to the image coordinate system, applies this correction, and stitches the sub-images into a single full-device image.

In the *Processing.ipynb* notebook, this is done automatically with the following function:

~~~
> stitcher.stitch_images()
~~~

The primary output from this step is a new directory of stitched images:

*experiments/*

*stitched_images*

Closely examine a stitched image. Good stitching should preserve the continuity of the chamber grid across tile boundaries. Some darkening near the edges of the original sub-images is expected, for example, due to optical vignetting. Still, chamber positions should remain continuous across tile borders (**Fig. 5B**). If tile boundaries are visibly discontinuous, the stitching should be repeated with adjusted rotation or stitching parameters before extracting fluorescence values.

#### Step 2. Subtract Background Images

During an experiment, for each type of image (kinetics assays, standard curves, or enzyme quantification) we take additional *background images*. These are images captured with the same exposure time, illumination intensity, optical filters, and camera settings as our assay images but without our fluorescent species of interest. These background images capture additive contributions from stray light, device and reagent autofluorescence, dust, and other sources of signal not attributable to enzyme or product fluorescence. *Mercury* uses the matched images to estimate and subtract this background signal from the corresponding assay data.

In the notebook, we specify our stitched background images and run the background subtraction cell:

~~~
>subtractor.subtract(…)
~~~

This automatically subtracts each background image, stored as a *NumPy* array (Harris et al., 2020), from any experimental image taken with the same camera settings.

The output of this step is a directory of background-subtracted stitched images:

*experiments/*

*bgsub_images*

#### Step 3. Segment Chambers and Buttons

Once stitched, background-subtracted images are generated; *Mercury then* segments the device into individual chambers and extracts RFU measurements. The HT-MEK device geometry provides a regular 32 × 56 chamber grid, corresponding to 1792 chambers. To align this grid to the image, the user specifies the image coordinates of representative corner chambers. *Mercury* then uses this grid to guide automatic feature finding, to identify device chambers and buttons using the Python package *scikit-image* (Walt et al., 2014).

Different image types are quantified using related but distinct segmentation strategies. For enzyme quantification, Mercury analyzes “button quant” images, in which the fluorescently tagged enzyme is localized to the button region of each chamber. The total fluorescence in the button region is quantified for each chamber and saved for later conversion into enzyme concentration, using the following function:

~~~
button_quant_processor.process()
~~~

For product standard curves and kinetic time courses, *Mercury* instead quantifies fluorescence within the chamber region, producing summed RFU values for each product concentration, substrate concentration, and time point:

~~~
standard_processor.process()
~~~

The latter notebook cells will calculate RFU values and save the output CSVs for downstream analysis. Expected output files at this step:

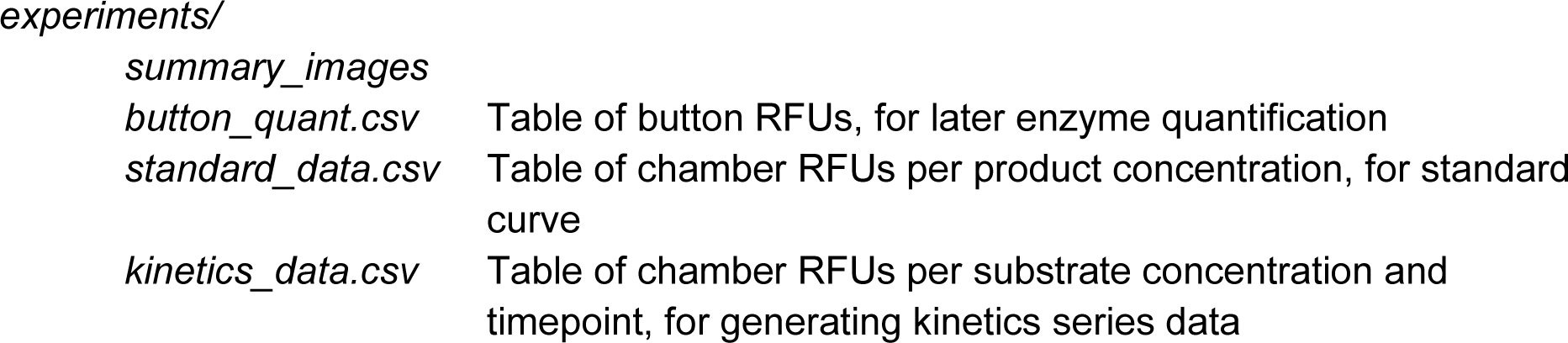

Segmentation quality should be evaluated using the summary images generated by *Mercury* (**Fig. 2C**). These images overlay the detected reaction-chamber and enzyme-immobilization (“button”) regions onto the processed image, allowing the user to confirm that the correct regions are being quantified. Representative summary images from across the device should be inspected to determine whether the segmentation masks align with the intended features and whether the expected features are visible in the underlying images. If the features are visible but the masks are shifted or missing, the grid-alignment or segmentation parameters should be adjusted and segmentation repeated. If the expected fluorescence is absent from the underlying image, the problem is upstream of segmentation and may reflect weak enzyme expression or immobilization, inadequate image acquisition, incomplete reagent delivery, or valve-actuation problems. Local segmentation failures should be reviewed against the underlying images.

### 4.3. Calibration of enzyme and product concentrations

After image processing, the *Mercury* workflow begins with three RFU tables: button eGFP fluorescence, product standard-curve fluorescence, and kinetic time-course fluorescence. These measurements are still reported in relative fluorescence units, so we begin by converting into experimentally meaningful quantities (i.e., concentrations) before rate fitting.

We first calculate the concentration of enzyme in each chamber using a single device-wide eGFP standard curve slope, which is pre-calculated on the microscope setup and used for all experiments. We *transform* our fluorescence data, dividing it by the slope of the eGFP standard curve. This is accomplished by the `> transform_data()` cell in the *Analysis.ipynb* notebook.

In the Jupyter notebook, a diagnostic plot shows enzyme concentrations across the device. Are these enzyme concentrations expected? Low concentration may indicate low expression during the experiment, due to poor plasmid purity, poor reagent flow during expression, or poor binding of the enzyme to the patterned nanobodies. Chambers with enzyme concentrations below our typical detection limit will be filtered at a later analysis step.

Next, we calculate the slope of the product standard curve in each chamber. As described in **Section 2.3**, per-chamber standard curves allow us to calibrate for anomalies that occur during the experiment and are calculated using the Python packages *SciPy* and *scikit-learn* (Pedregosa et al., 2011; Virtanen et al., 2020).

~~~
> standard_fits = fit_Lumi.nance_vs_concentration(…)
~~~

In the diagnostic plot, note the regular brightening and darkening pattern across the device, corresponding to higher or lower standard curve slopes. This is expected and results from optical vignetting, among other factors. Our per-chamber calculated standard curves will correctly account for this effect. Examine your data -are there wells with atypical standard curve slopes? Because the standard curve is generated by flowing product directly into the device, anomalies that run vertically in columns likely indicate poor solution flow on the device. Alternatively, single outlier chambers may indicate debris obscuring imaging. These chambers will be filtered in subsequent steps.

After fitting our standard curve slopes, we convert our summed RFU values for our kinetics series data into units of product concentration using a similar notebook cell analogous to the enzyme-concentration calibration described above. At the end of this step, *Mercury* has generated per-chamber enzyme concentration and product-concentration time courses. These calibrated data form the input to initial-rate fitting.

### 4.4. Initial-rate Fitting and Background Rate Subtraction

We now fit initial rates to our reaction progress curves. Because this is a crucial and nuanced step in our analysis, refer to **Section 3.6** for detailed considerations in selecting fitting windows.

~~~
kinetics_fits, … = fit_concentration_vs_time(…)
~~~

Examine the diagnostic plot; clicking on a chamber reveals the initial rate fits for that chamber. Do the linear fits appear to capture the slope of the initial phase of the reaction? Follow the prompts in the notebook to tweak starting and ending time points, either globally or per concentration, if necessary.

Because many reactions have an appreciable background rate in solution, we aim to separate this effect from our enzyme-catalyzed rates. In the next cell (not pictured), we select chambers that contain no protein, or a non-catalytic control (like eGFP). *Mercury* estimates the background rate from these control chambers (**Section 2.4**) and subtracts it from the measured initial rates before downstream fitting.

After fitting and background rate subtraction, *Mercury* saves per-chamber and per-concentration slopes, intercepts, and R^2^ values for the fits.

### 4.5. Quality Control: Filtering and Flagging Chambers and Initial Rates

At this point in the workflow, *Mercury* has calculated enzyme concentration, product standard-curve parameters, product concentration time courses, and background-corrected initial rates. The next task is to decide which measurements are reliable enough to pass into Michaelis-Menten fitting.

*Mercury* performs filtering by generating masks: Boolean arrays that indicate which chambers or measurements should be carried forward into downstream analyses. Previewing these masks serves as both a quality-control and diagnostic step. While individual chambers can be excluded if they do not meet a given criterion (e.g., enzyme expression is below our limit of detection), large-scale flagging of chambers suggests an issue with upstream analysis that should be examined carefully. The most common filters are summarized in **Table 3**.

**Table 3:**
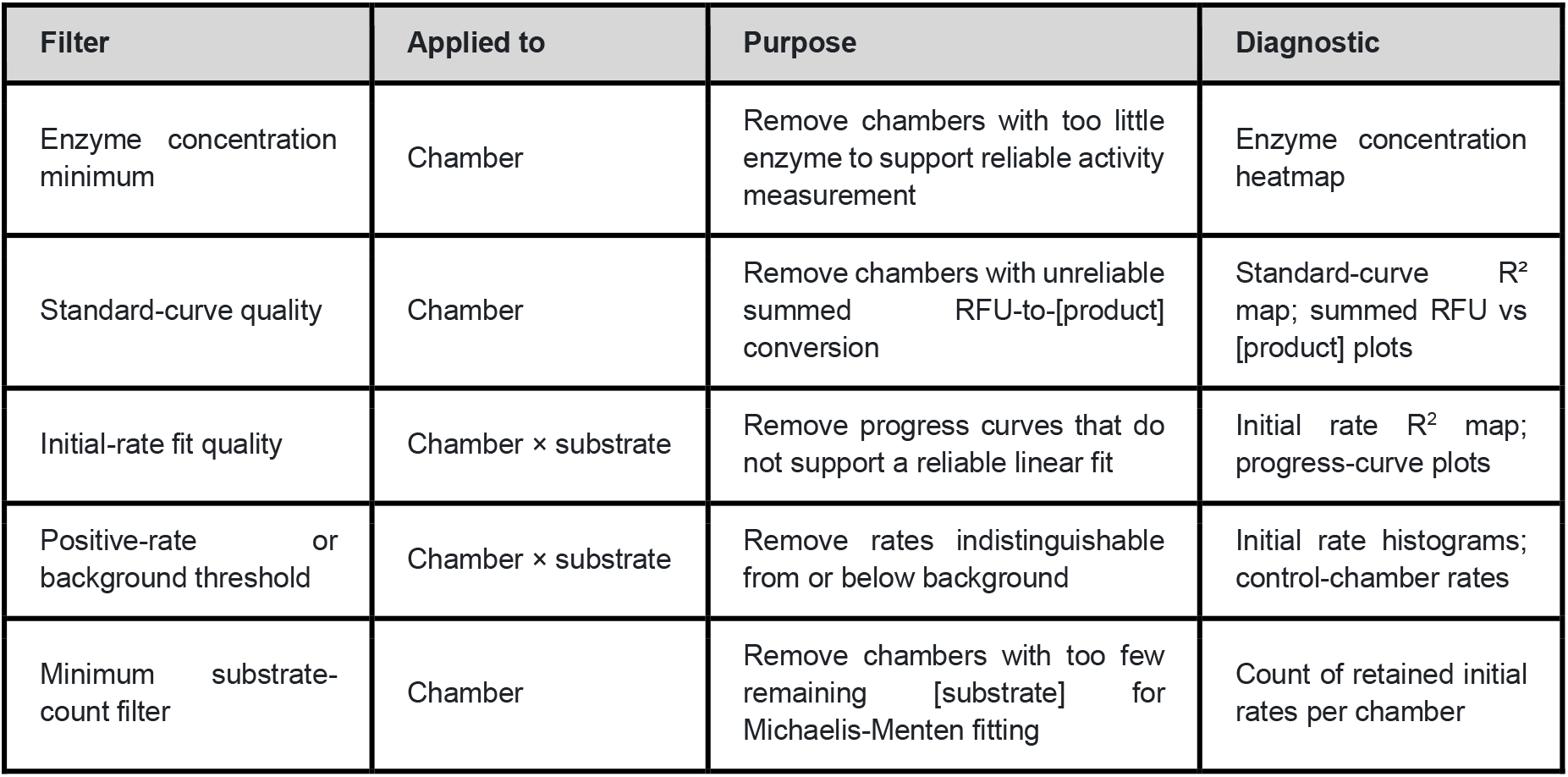
Filters applied to initial-rate fits.

After individual filters have been defined, *Mercury* combines them into a final mask and applies the mask to the initial-rate data. The number and spatial distribution of retained measurements should then be inspected. If many chambers have too few remaining substrate concentrations, Michaelis-Menten parameters will not be well constrained. In the workflow described here, chambers must retain at least 5 substrate concentrations before Michaelis-Menten fitting is attempted. For reproducibility and documentation, ensure you save the filtering criterion used: we recommend saving a copy of the *Analysis.ipynb* notebook in its entirety. At the end of this step, *Mercury* has saved filtered copies of our initial-rate data.

### 4.6. Fitting Michaelis-Menten parameters

After flagging and filtering, the retained initial rates are used to fit Michaelis-Menten parameters using the following function:

~~~
MM_fits, … = fit = fit_initial_rates_vs_concentration_with_function(…)
~~~

We fit the Michaelis-Menten equation (**Section 2.1**) to the data from each chamber, yielding V_max_, *K*_M_, and the fit R^2^. Dividing each chamber’s V_max_ by its enzyme concentration yields *k*_cat_.

Once chamber-level fits have been generated, *Mercury* aggregates replicate chambers belonging to the same sample. For each sample, the retained chamber-level estimates are combined to report sample-level kinetic parameters and replicate variability. Next, we utilize z-score filtering to flag chambers for visual inspection. The resulting output includes the mean and standard deviation of the fitted parameters, fit-quality statistics, and the number of replicates retained after filtering.

### 4.7. Exporting results and diagnostics

In this final step, we export fit catalytic parameters and statistics. While we are interested in the aggregate data for each sample, we also export per-chamber data for diagnostics.

Additionally, we generate several useful plots to understand and diagnose potential issues with data. Final outputs from *Mercury* analysis are shown in **Table 4**.

**Table 4.** Outputs from *Analysis.ipynb*.

| Output | Level | Purpose |
| --- | --- | --- |
| MM_sample_data.csv | Sample | Main quantitative output for downstream analysis. Includes kinetic parameters and relevant statistics. |
| MM_chamber_data.csv | Chamber | Diagnostic table of chamber-level fits and filtering status |
| MM_sample_subplots.pdf | Sample | Michaelis-Menten plots for visual inspection of fitted parameters |
| summary_plots/ | Selected samples | End-to-end diagnostic summaries linking image quality to final fits |
| Saved notebooks/configuration | Analysis | Record of fitting windows, filters, and calibration choices. |

The data in *MM_sample_data.csv* serves as the primary output, to be used in downstream analysis, such as relating enzyme catalytic parameters to structural features or comparing ortholog activity across a phylogenetic tree. The other generated files provide varied diagnostics. In particular, the sample-level summary plots (**Fig. 7**) integrate raw experimental images with their corresponding analyzed plots to provide a holistic understanding of the input data quality, model fitting, and final output accuracy. These visual summaries offer additional guidance for verifying data integrity and checking the validity of kinetic parameter fits. *Note: Because compiling the complete set of summary plots can require up to 30 minutes, beginning with a specific subset of interest is highly recommended*.

**Figure 7.**
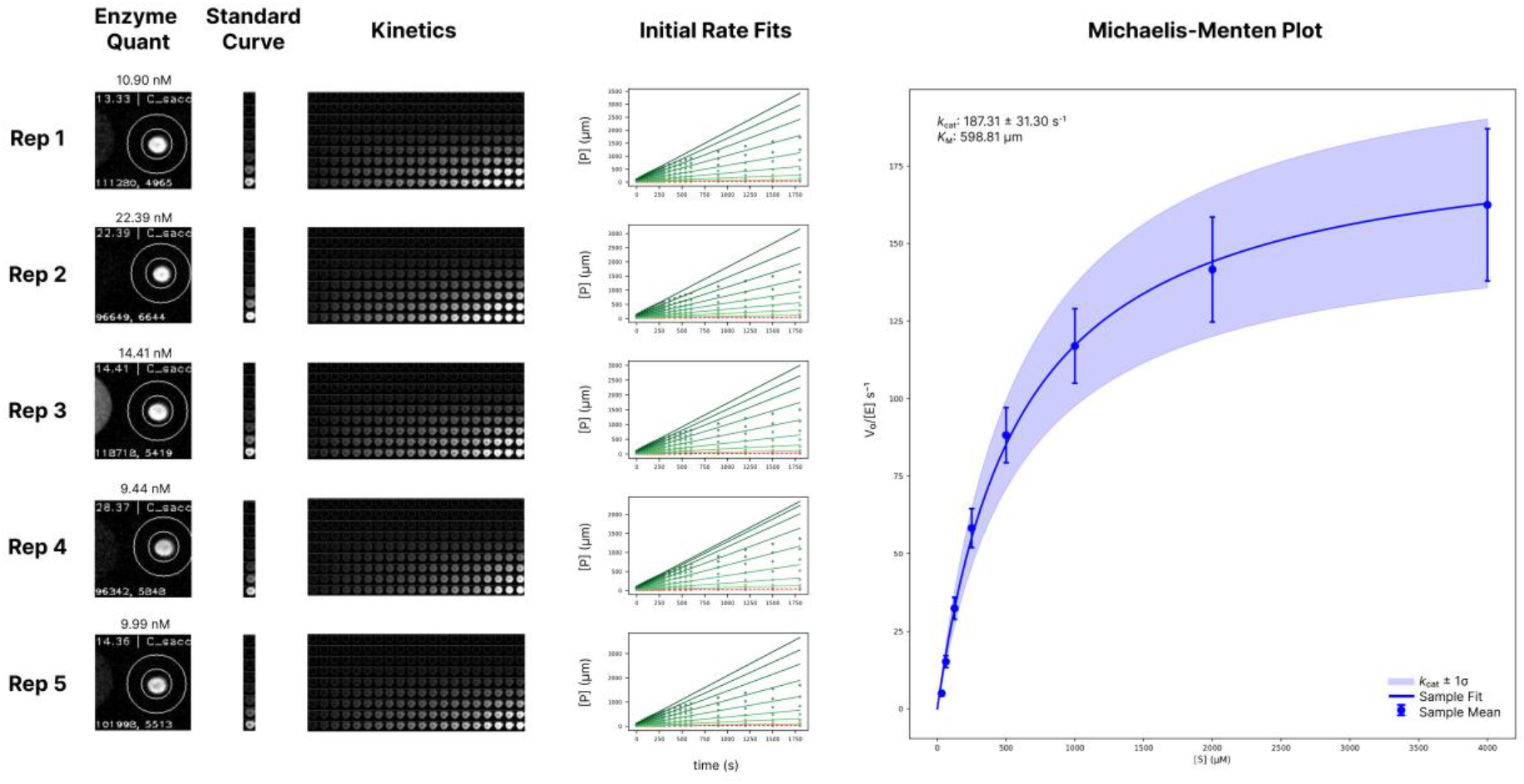
End-to-end summary plot of one sample after *Mercury* Analysis. This plot synthesizes experimental images with analysis plots for human review. Per-chamber images of enzyme quantification, standard curve, and reaction kinetics are displayed to reveal possible sources of error. Reaction progress curves are shown, with fit initial rates plotted. Datapoints or initial rates that have been excluded from downstream analysis during data filtering are shown in red with dashed fit lines. The final Michaelis-Menten plot aggregates replicate Michaelis-Menten fits into a single per-sample plot, with sample means and fit envelope displayed. Fonts and spacing have been adjusted for publication.

## 5. Data and code availability

*Mercury* is open-source and released under the permissive MIT License. Code, installation instructions, and documentation are available at: https://github.com/pinneylab/mercury/.

Jupyter notebooks and example datasets are available at: https://github.com/pinneylab/mercury_notebooks/.

## 6. Conclusion

High-throughput enzymology increases the scale at which relationships between enzyme sequence and kinetics can be dissected. Nevertheless, the value of this scale depends on maintaining a clear connection between reported parameters and the measurements from which they were derived. Thus, the underlying analytical workflows should preserve this connection by documenting data transformations, model assumptions, quality-control criteria, replicate-level results, and parameter uncertainty. For processing and analyzing HT-MEK data, diagnostic visualization is central because it allows numerical outputs to be evaluated in the context of the underlying raw and processed images, progress curves, calibrations, and controls.

Although the workflow presented here uses Michaelis-Menten analysis of HT-MEK data as its primary example, these principles apply to a broad range of quantitative biochemical assays. *Mercury*’s modular organization allows this workflow to be extended to binding, inhibition, and other biophysical measurements which are already being pioneered in the HT-MEK and related platforms (Aditham et al., 2021; Markin et al., 2023), although each application requires assay-specific models, controls, and quality-control criteria. As high-throughput biochemical datasets continue to expand, transparent documentation of data processing and accessible diagnostic information will facilitate comparison, reanalysis, and methodological refinement.

## 7. Glossary of Terms

Background image: An image acquired with the same microscope and camera settings as an assay image, but without the fluorescent species of interest. It captures camera offset, stray light, device autofluorescence, dust, and other off-target signals.
Background rate: The reaction rate measured in no-enzyme or non-catalytic control chambers, representing non-enzymatic product formation and other rate-like background signals. In the workflow described here, it is estimated separately at each substrate concentration from the lower quartile of the control-chamber rates.
Background-image subtraction: Pixel-wise subtraction of a background image from a corresponding assay image to remove fluorescence not attributable to the enzyme or product being measured.
Background-rate subtraction: Subtraction of the estimated background rate from the measured initial rate of each sample at the corresponding substrate concentration. This is distinct from image background subtraction.
Button: A circular, pneumatically actuated membrane within an HT-MEK unit cell. The button can protect or expose a defined surface region during enzyme immobilization, washing, and reaction setup. Fluorescence within the button region is used to quantify immobilized eGFP-tagged enzyme.
eGFP tag: An enhanced green fluorescent protein fused to the enzyme of interest. The tag provides a fluorescence signal for estimating enzyme concentration and enables enzyme localization through capture by an anti-eGFP nanobody.
High-Throughput Microfluidic Enzyme Kinetics (HT-MEK): A microfluidic platform for performing parallel biochemical measurements across many enzyme variants, conditions, and replicates. HT-MEK independently estimates enzyme concentration and monitors reaction progress across substrate concentrations, allowing catalytic parameters to be determined for individual variants.
Image segmentation: Identification of the chamber, button, or other device regions from which fluorescence measurements will be extracted.
Initial rate (V0): The slope of the early, approximately linear portion of a product-concentration-versus-time curve. It estimates the reaction rate before substantial substrate depletion, product inhibition, reverse reaction, or enzyme inactivation occurs.
Initial-rate fitting window: The subset of time points used to fit the line from which (v_0) is calculated. The window must be long enough to determine a precise slope but short enough to remain within the initial linear regime.
Kinetic regimes: Regions of a substrate titration with different parameter information. In the low-substrate regime, enzyme-normalized rate is approximately linear with substrate concentration and primarily constrains *k*_cat_/*K*_M_. In the saturating regime, the rate approaches a plateau and primarily constrains V_max_ and *k*_cat_.
Mask: A Boolean array indicating which chambers or measurements should be retained and which should be excluded. Mercury combines individual quality-control masks into a final mask before downstream fitting.
Mercury: The open-source Python analysis framework described in this chapter. Mercury processes HT-MEK microscopy images, performs fluorescence calibration, fits initial rates and kinetic models, applies quality-control filters, aggregates replicates, generates diagnostic visualizations, and exports results. It also supports user-defined models and assay-specific workflows.
Optical vignetting: Spatial variation in optical sensitivity across an image, typically producing lower sensitivity near the image edges than near its center. HT-MEK analysis compensates for this effect using a separate product standard curve for each chamber.
Per-chamber product standard curve: A calibration relating known product concentrations to fluorescence within one reaction chamber. Separate standard curves are fitted for each chamber to account for vignetting and other spatial differences in illumination, detection, and background fluorescence.
Reaction chamber: An individual microfluidic compartment in which an enzyme is immobilized, a reaction is isolated, and fluorescence is measured. The HT-MEK device described here contains a regular array of 1,792 reaction chambers.
Reaction progress curve: A time course showing measured or inferred product concentration as a function of time for a particular chamber and substrate concentration.
Refit, filter, qualify, or repeat: The four principal responses in the chapter’s quality-control framework. Refit changes an inappropriate analysis choice; filter excludes localized invalid measurements; qualify reports conclusions with limitations when only some parameters are supported; and repeat is used when an upstream or global failure cannot be corrected analytically.
Relative fluorescence units (RFU): Arbitrary units representing measured fluorescence intensity. In this workflow, RFU generally refers to the summed pixel intensity within a segmented chamber or button region.
Replicate chamber: A separate reaction chamber containing the same enzyme variant or sample as another chamber. Replicates are distributed across the device to reduce the risk that a spatially patterned failure affects all replicates of a sample.
Scooping: Early upward curvature in a reaction progress curve before a stable linear regime is reached. It may indicate incomplete mixing, delayed reaction initiation, or valve-timing effects.
Spatial map or heatmap: A visualization in which chamber-level measurements are arranged according to their physical positions on the microfluidic device. These plots help distinguish isolated failures from rows, columns, gradients, tile boundaries, and other spatial patterns.
Stitching: Rotation, alignment, and merging of partially overlapping microscopy tiles into a single contiguous image of the full HT-MEK device.
Substrate titration: A series of measurements collected across multiple substrate concentrations and used to fit a kinetic model or determine which kinetic parameters are supported by the data.
Summary image: A diagnostic processed image that overlays the chamber or button regions detected by Mercury on the microscopy image, allowing the user to inspect segmentation and signal quality.
Tile or sub-image: An individual microscopy image covering only part of the HT-MEK device. Multiple overlapping tiles are acquired at each condition or time point and subsequently stitched together.
Valve: A pressure-controlled microfluidic feature that directs fluid movement and isolates reaction chambers during enzyme immobilization, washing, reagent loading, and reaction measurement.

## 8. Acknowledgments

We thank members of the Pinney lab for helpful discussions and critical feedback on the manuscript, for testing the Mercury workflow and documentation, and for contributions of experimental images. We also acknowledge the foundational development of the HT-MEK experimental and analytical framework by members of the Fordyce laboratory. M.M.P. is a Biohub, San Francisco, Investigator.

## 9. Declaration of AI and AI-assisted Technologies in the Writing Process

During the preparation of this work the authors used ChatGPT and Gemini in order to improve word choice and clarity of sections of the manuscript. ChatGPT was used to generate a microscope illustration in Fig. 3d. After using these tools/services, the authors reviewed and edited the content as needed and take full responsibility for the content of the publication.

